# Chemical Gating of the Mechanosensitive Piezo1 Channel by Asymmetric Binding of its Agonist Yoda1

**DOI:** 10.1101/169516

**Authors:** Jerome J. Lacroix, Wesley M. Botello-Smith, Yun Luo

**Author notes:** corresponding author: Dr. Jerome Lacroix Western University of Health Sciences 309 E. Second St, Pomona, CA 91766.

## Abstract

Piezo proteins are homotrimeric ion channels that play major roles in normal and pathological mechanotransduction signaling in mammalian organisms. Their pharmacological control hence represents a potential therapeutic avenue. Yoda1, a Piezo1-selective small molecule agonist, is the only known selective Piezo modulator. How Yoda1 selectively interacts with Piezo1 and opens its pore is unknown. Here, by engineering and characterizing chimeras, we identified a minimal region responsible for Yoda1 binding. This region is located at the interface between the pore and the putative mechanosensory domains in each subunit. By characterizing hybrid channels containing Yoda1-insensitive and Yoda1-sensitive monomers, we demonstrate that the presence of only one Yoda1-sensitive Piezo1 subunit is sufficient for chemical activation, implicating that the asymmetric binding of Yoda1 to a single subunit enables channel opening. These findings shed light onto the gating mechanisms of Piezo channels and will pave the way for the rationale design of new Piezo channels modulators.

## Introduction

Piezo proteins are very large (~1MDa) trimeric mechanosensitive ion channels (Fig 1a) which transduce various forms of mechanical stimuli such as shear stress and membrane stretch into important biological signals. In mammals, only two Piezo isoforms named Piezo1-2 have been identified. In spite of their recent discovery, these isoforms have been implicated in a bewildering number of mechanotransduction processes including touch sensation(1–3), proprioception(4), hearing(5), vascular(6, 7) and brain development(8), blood flow sensing(9), osmotic homeostasis(10) and epithelium regulation(11, 12). Both gain-of-functions and loss-of-functions Piezo mutations are associated with human diseases such as xerocytosis(13, 14, 10, 15), arthrogryposis(16–22) and lymphedema(23). Recent studies suggest Piezo channels may also play important roles in other conditions such as sleep apnea(24) and visceral pain(25).

**Fig. 1.**
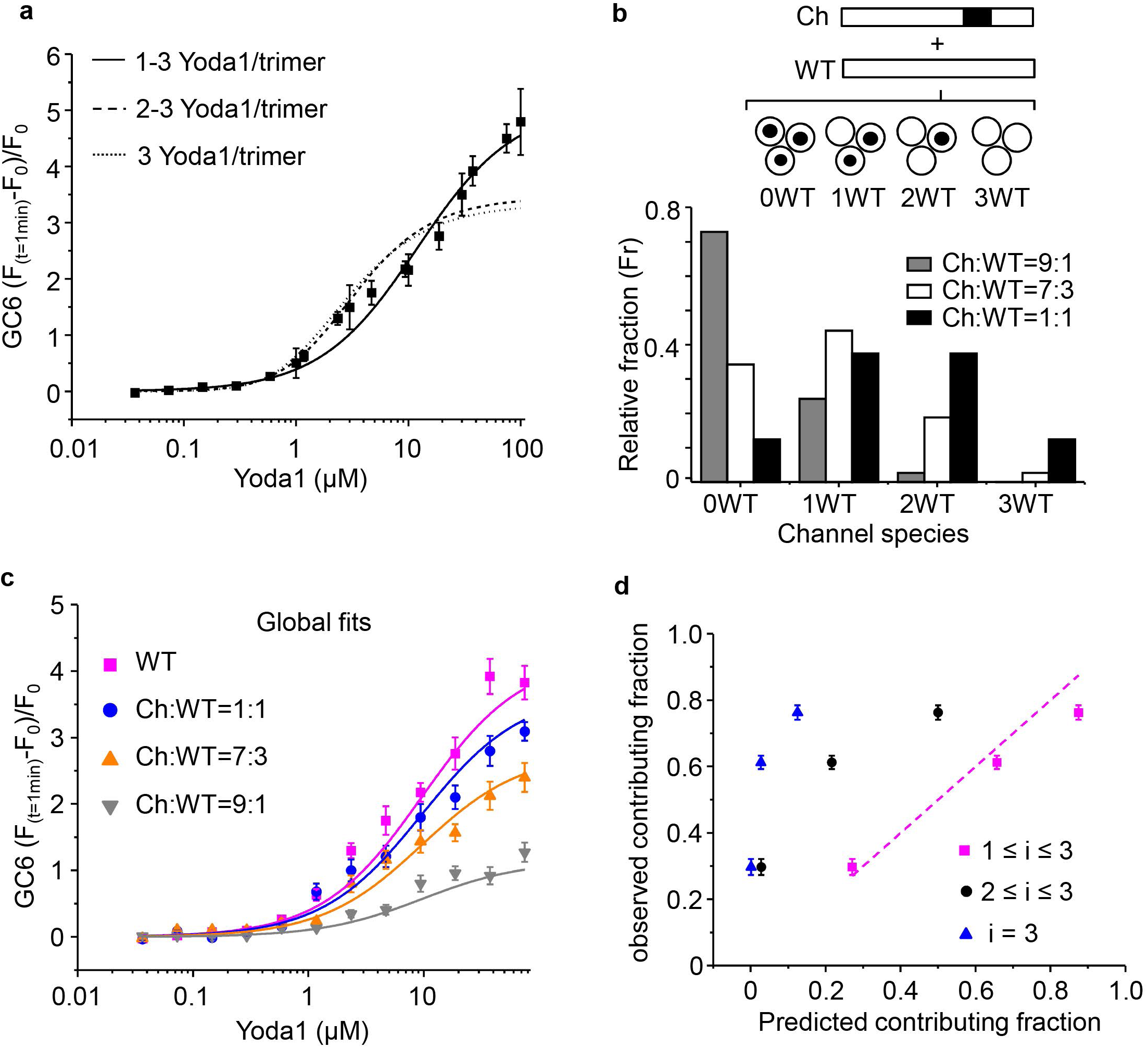
A chimeric approach to determine the Yoda1 binding site in mPiezo1. **a** mPiezo1 cryo-EM structure (PDBID:3JAC). The indicated structural features are highlighted with differential coloring. **b** Yoda1 chemical structure. **c** Generation of C-terminal chimeras with respect to their approximate position in the mPiezo1 topology (coloring is identical to a). Residue numbers are from mPiezo1; mPZ1=mPiezo1, mPZ1=mPiezo2. **d** Relative Ca^2+^-sensitive fluorescence time course from HEK293T cells transfected with WT mPZ1 (black and magenta traces) or the empty vector pCDNA3 (blue trace). Cells were pre-loaded with Fluo8-AM and incubated with 30μM Yoda1 (blue and black traces) or a control solution (magenta trace) at t = 30 s. **e** Relative fluorescence changes from HEK293T cells expressing the indicated C-terminal chimeras (n = 4 for each plot), mPZ1 (magenta, n = 4) or mPZ2 (purple, n = 4) plotted against Yoda1 concentration at t = 60 s. **f** Relative fluorescence changes from HEK293T cells expressing the indicated construct (n = 5 for each plot) or the empty vector pCDNA3 (control, n = 5) following an acute hypotonic shock. For each construct, the fluorescence signal was compared to the control using a Student’s t-test. The numbers of the left of the bars above the histograms indicate the t-values for each test. Asterisks indicate standard p-value range: ^*^: 0.01 < p < 0.05; ^**^: 0.001 < p < 0.01 and ^***^: p < 0.001. In all panels, error bars = s.e.m.

An avenue to treat these diseases would be to correct Piezo channel activity with small molecule agonists and/or antagonists. On the other hand, the availability of selective Piezo modulators would allow us to precisely determine the contribution of each Piezo isoform to complex physiological functions. Unfortunately, the current pharmacology of Piezo channels is extremely limited, with a single Piezo1-selective small molecule agonist named Yoda1(26) (Fig 1b) and no known isoform-selective inhibitors. Hence, understanding how Yoda1 selectively binds and activates Piezo1 is a key step to rationally design new small molecule modulators with high selectivity.

Yoda1 was recently identified using a simultaneous high-throughput agonist screening on both Piezo1 and 2 isoforms. Although Piezo2 shares more than 65% of its primary amino acid sequence with Piezo1, Yoda1 possesses strict Piezo1 selectivity with no measurable modulatory effects on Piezo2 activity. The activation of Piezo1 by Yoda1 originates from the binding of Yoda1 to one or more unknown binding site(s) on the Piezo1 channel(26). Based on these observations, we have generated Piezo1-Piezo2 chimeras to identify a minimal protein region responsible for Yoda1 binding selectivity. By characterizing hybrid channels containing wild-type (WT) and Yoda1-insensitive Piezo1 subunits, we further show that the binding of Yoda1 to a single subunit is sufficient for activation of the trimeric channel.

## Results

### Molecular determinants for Yoda1 selectivity

The purified Piezo1 channel reconstituted in artificial bilayer remains Yoda1-sensitive(26). Hence, the Piezo1 protein possesses an endogenous agonist Yoda1 binding site that enables channel activation. The strict Piezo1-selectivity exhibited by Yoda1 must then originate from amino acid differences between the Piezo1 and Piezo2 primary sequence. Since chimeras between evolutionary-distant Piezo homologs remain functional(27), we reasoned that chimeras between mouse Piezo1 (mPiezo1) and mouse Piezo2 (mPiezo2) will yield functioning proteins with altered Yoda1-sensitivity. Since Yoda1 stabilize the open pore conformation in absence of mechanical stimulation(26), we reasoned that the Yoda1 binding site is likely located near the pore in the C-terminal region. We thus designed three C-terminal chimeras and named them by the residue number after which the C-terminal sequence is from mPiezo2 (i.e. Ch1961, Ch2063 and Ch2456, Fig 1c and Supplementary Fig 1). The C-terminal chimeras were expressed into HEK293T cells and characterized using a fluorescence assay based on the calcium-sensitive organic dye Fluo8-AM (Fig 1d). A variant of this assay was previously used to characterize Yoda1 sensitivity of Piezo homologs and disease variants(26). The chimeras Ch2063 and 2456 exhibit concentration-dependent fluorescence responses similar to WT mPiezo1 (Fig 1e). In contrast, Ch1961 was totally insensitive to Yoda1 in the range of concentrations tested, similar to the Yoda1-insensitive mPiezo2. We independently confirmed the ability of the three C-terminal chimeras to respond to mechanical stimulation induced by an acute hypotonic shock, a simple way to experimentally activate Piezo channels. All chimeras displayed robust responses similar to mPiezo1. These responses were significantly larger than the response obtained from cells transfected with a control vector (Fig 1f).

### A minimal region for Yoda1 binding

To verify that the region 1961-2063 contains the molecular determinants for Yoda1 binding, we created the internal chimera Ch1961-2063. We next divided this region into three sub-regions (region 1: 1961-2004, region 2: 2005-2034 and region 3: 2035-2063) and engineered every possible combination of “sub-chimeras” replacing one or two sub-regions by the corresponding sequence from the mPiezo2 homolog (Fig 2a and Supplementary Fig 1).

**Fig. 2.**
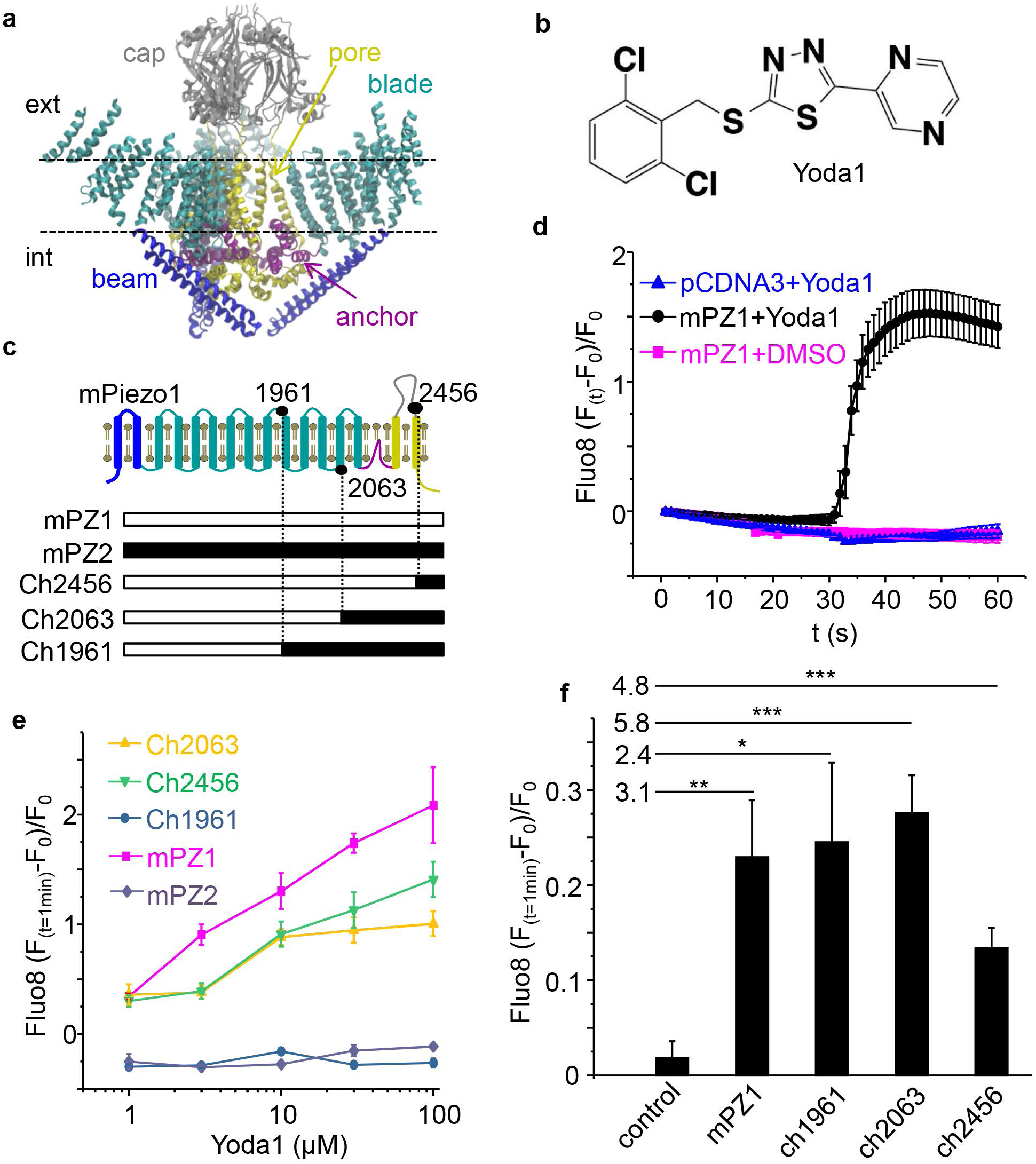
A minimal mPiezo1 region responsible for Yoda1 binding. **a** Generation of internal chimeras in the mPiezo1 1961-2063 region. Filled regions represent mPZ2 sequence; residue numbers are from mPiezo1. **b** Relative fluorescence changes from HEK293 cells expressing the indicated Chimeras (n = 4 for each plot), mPZ1 (magenta, n = 4) or mPZ2 (dark blue, n = 4) plotted against Yoda1 concentration. **c** Relative fluorescence changes from HEK293T cells expressing mPZ1 (n = 6), mPZ2 (n = 5) the Ch1961-2063 chimera (n = 6) or the empty vector pCDNA3 (control, n = 6) following an acute hypotonic shock. For each construct, the fluorescence signal was compared to the control using a Student’s t-test. The numbers of the left of the bars above the histograms indicate the t-values for each test. Asterisks indicate standard p-value range: ^*^: 0.01 < p < 0.05; ^**^: 0.001 < p < 0.01 and ^***^: p < 0.001. **d** Pressure-clamp electrophysiology recordings in the cell-attached configuration for mPZ1 (magenta traces) and the Ch1961-2063 chimera (blue traces). **e** Normalized peak current as a function of the pressure pulse for mPZ1 (magenta squares, n = 3) and Ch1961-2063 (blue circles, n = 3). The traces correspond to a fit using a classical two-state Boltzmann equation (see text). **f** Position of the 1961-2063 sub-regions (1961-2004, yellow; 2005-2034, orange and 2035-2063, blue) relative to the mPiezo1 cryo-EM structure (left: top view, right: side view). In all panels, error bars = s.e.m.

For simplicity, these sub-chimeras were named by the number of the sub-region(s) being replaced (e.g. Ch1+3 in Fig 2a replaces the sub-regions 1961-2004 and 2035-2063 by their homolog sequences from mPiezo2). To reduce fluorescence background, calcium-sensitive fluorescence was measured by co-transcriptionally expressing the genetically-encoded fluorescent indicator GCaMP6m (see Methods). Wide-field fluorescence live-cell imaging showing GCaMP6m time course upon Yoda1 application in the presence or absence of mPiezo1 can be seen in Video1 and Video2, respectively. The fluorescence of GCaMP6m does not change upon Yoda1 application in absence of mPiezo1, as when using Fluo8-AM (Fig 1d). All tested sub-chimeras displayed diminished fluorescence responses compared to mPiezo1 (Fig 2b). Yet, none of them completely suppressed the fluorescence response as seen in cells transfected with mPiezo2 or with the internal chimera 1961-2063. The internal chimera 1961-2063 remains able to elicit robust fluorescence responses upon acute hypotonic chock (Fig 2c) and to produce mechano-dependent ionic currents with amplitude similar to wild type mPiezo1 (Fig 2d). This shows that the elimination of the fluorescence signal in cells transfected with Ch1961-2063 originates from the loss of the agonist binding site rather than a loss of channel function. We next fitted the plots of the normalized peak currents I/I_max_ against the pulse pressure P (Fig 2e) with a standard two-state Boltzmann equation:

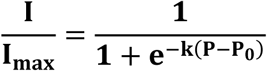

with P_0_ the pressure corresponding to I = I_max_/2 and k a constant representing the intrinsic electromechanical coupling of the channel. The fits yielded P_0_ = 32.29 ± 0.93 mmHg for WT mPiezo1, consistent with previous studies(26). However, the chimera Ch1961-2063 exhibits an increased threshold for mechanosensitivity with a fitted P_0_ value of 56.43 ± 0.76 mmHg (see Supplementary Table 1).

### Minimal Yoda1-sensitive subunit stoichiometry for Yoda1-mediated activation

The presence of the entire mPiezo1 1961-2063 region appears necessary for chemical activation with Yoda1. Interestingly, this region is strategically located at the interface between the pore and the blade in each subunit (Fig 1a and Fig 2f). Since this region exists in each of the three Piezo1 subunits, Yoda1 could potentially bind to each subunit. How many subunits need to interact with the Yoda1 agonist to open the channel’s pore?

In our assay, the measured fluorescence F is a function of the saturation function v of the protein P by the ligand L and of the background fluorescence q:

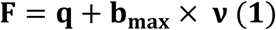

At t = 0 s, [Yoda1] = 0 μM, then ν =o:

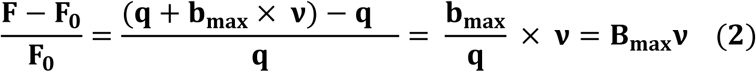

In case of multiple binding sites per protein, the saturation function equals the total concentrations of bound ligands ([L]_bound_) over the total concentration of protein ([P]_0_):

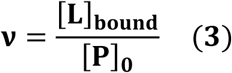

Assuming the existence of three identical and independent Yoda1 binding sites:

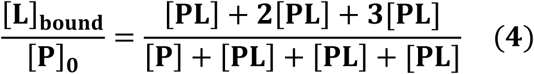

The different protein species are related by the corresponding macroscopic dissociation constant K_1_, K_2_ and K_3_ corresponding to each binding step:

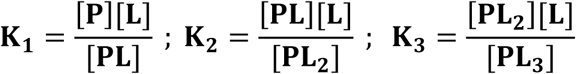

By replacing the macroscopic constants

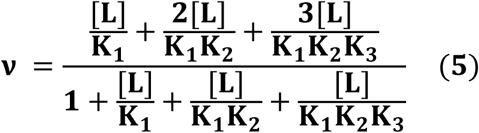

The macroscopic constants are related to the microscopic dissociation constant K_d_ by the numbe or possible binding combinations Ω_n,i_ for each binding step i and for n binding sites:

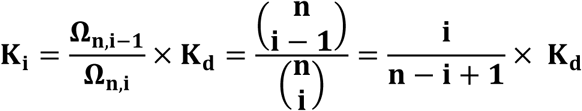

Replacing the macroscopic constants by the Kd gives:

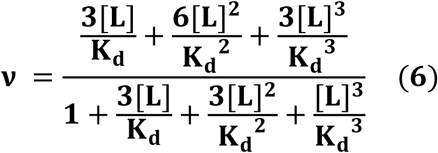

Equation (6) further simplifies by applying a binomial reduction:

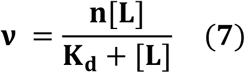

The details about this reduction can be found elsewhere(28). Assuming the binding of one or more ligand per channel produces similar channel opening, every bound species in the numerator of equation (6) contribute proportionally to the observed fluorescence signal:

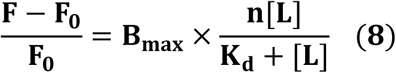

However, if the binding of two or three ligands is required for channel activation, the fraction of channel with a single ligand does not contribute to the signal. In this case, the fluorescence signal follows:

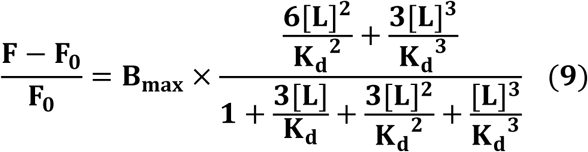

Similarly, if the binding of three ligands is necessary for channel activation, the fluorescence signal follows:

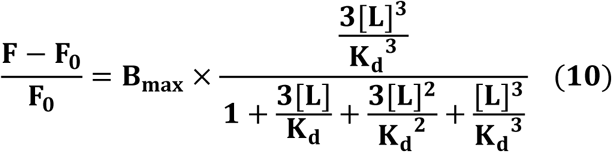

We tested these three binding situations modeled by equations (8), (9) and (10) by performing a dose-response on WT mPiezo1 with more data points (Fig 3a). Curve fitting clearly shows that a model with 3 ligands per channel (equation (10)) or with more than 2 ligands (equation (9)) does not fit well the data (Fig 3a and Table 1). In contrast, a model where every bound species equally contribute to the signal (equation (8)) provides a better fit. This suggests the binding of Yoda1 to a single binding site in one subunit is sufficient for channel opening. As indicated previously, the reported Kd values obtained from curve fitting are not accurate estimates of the dissociation constant due to the poor aqueous solubility of Yoda1 above 20-30 μM(26).

**Fig. 3.**
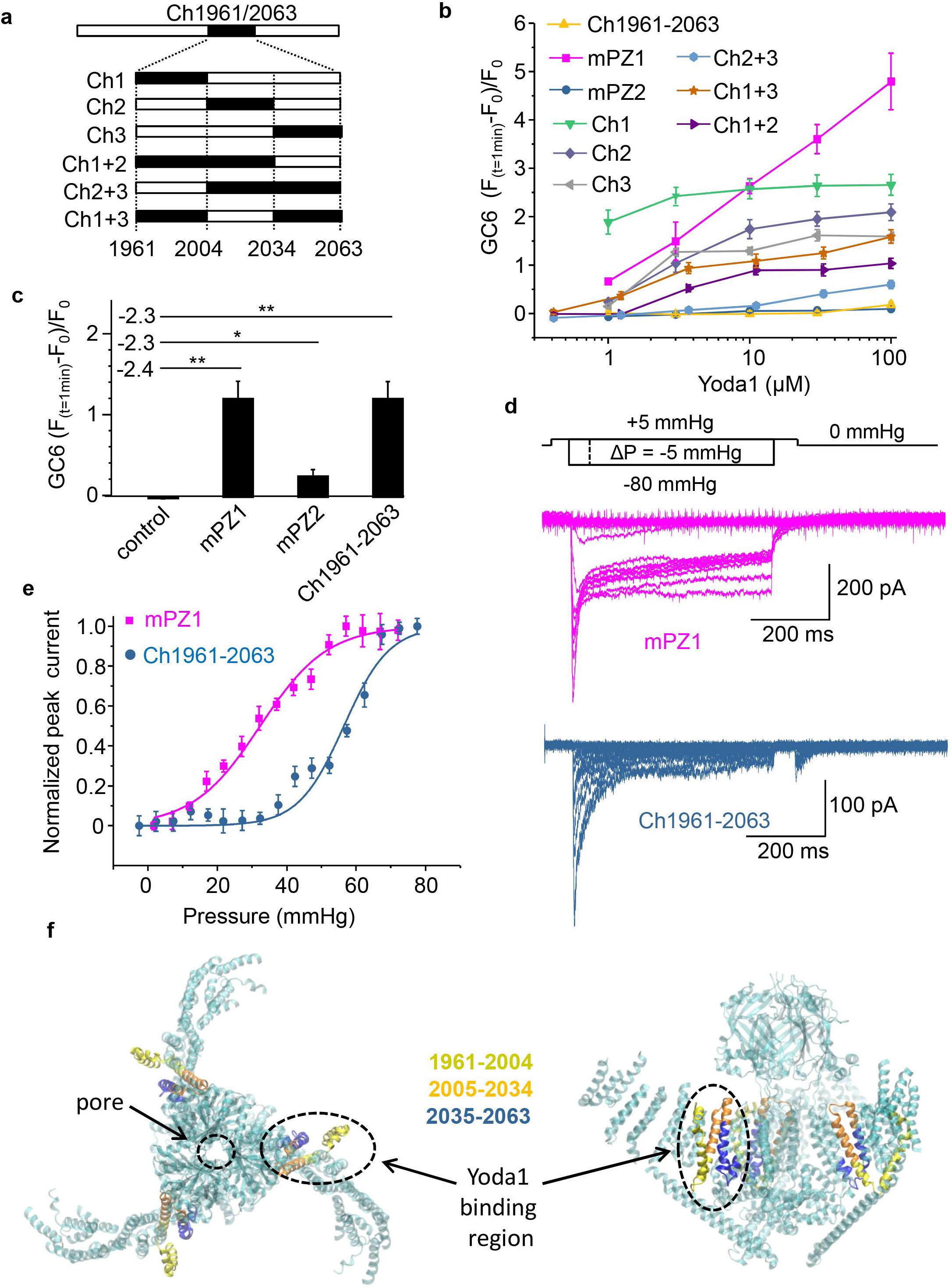
Minimal effective Yoda1 binding stoichiometry. **a** Fitting of the mPiezo1 Yoda1 dose response (n = 4) with three binding models corresponding to equations (8), (9) and (10) (see text and Table 1 for fitted parameters). **b** Co-expression of WT mPiezo1 subunits (WT) with Yoda1-insensitive Ch1961-2063 chimera subunits (Ch) yields four distinct channel species with predicted relative fraction as a function of the relative amount of WT vs. Ch subunits. **c** Yoda1 dose-responses obtained by mixing WT and Ch subunits with the indicated ratios (n= 4 for each plot). The traces are fit obtained with a global fit from equation (12). **d** Observed vs. predicted fraction of contributing hybrid channels having i WT subunit. Each point is the mean values for six concentrations ranging from 2 to 75 μM for the three tested mixtures. The red line corresponds to the linear function y = x. In all panels, error bars = s.e.m.

**Table 1:** Fitting results for different binding stoichiometry

| Number of ligand per trimeric channel | 1-3 | 2-3 | 3 |
| --- | --- | --- | --- |
| Model equation | (8) | (9) | (10) |
| $B_{\max}$ | $1.701 \pm 0.152$ | $1.155 \pm 0.097$ | $0.744 \pm 0.102$ |
| $K_d$ ($\mu\text{M}$) | $11.990 \pm 1.941$ | $2.245 \pm 0.297$ | $1.110 \pm 0.103$ |
| $R^2$ | 0.964 | 0.812 | 0.825 |

To further investigate the minimal binding stoichiometry of Yoda1, we co-expressed WT mPiezo1 subunits with Yoda1-insensitive Ch1961-2063 subunits by transfecting HEK293T cells with different ratios of the corresponding plasmids. The subunit mixtures lead to four channel species with a number of WT subunits ranging from 0 to 3 (Fig 3b). Assuming WT and chimeric subunits are expressed in proportion to the amount of transfected plasmid and assuming hybrid channels are formed by random association of WT and chimeric subunits, the fraction (Fr) of assembled channels with *i* WT subunits for a trimeric channels is given by the equation:

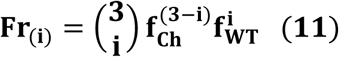

With f_Ch_ and f_WT_ the relative fractions of plasmids encoding chimeric and WT subunits, respectively.

We transfected HEK293T cells with different plasmid mixtures containing 10%, 30%, 50% or 100% WT mPiezo1. The predicted fraction for each species and for each tested mixture is indicated in Fig 3b. Surprisingly, cells transfected with a mixture containing only 10% WT subunits produced robust fluorescence signals (Fig 3c). Indeed, at saturating Yoda concentrations, this signal amounts to approximately a third of the fluorescence signal observed in cells transfected with 100% WT plasmids. When mixing 10% WT subunit with 90% chimeric subunit, the predicted fraction of channels containing three WT subunits is only 0.1%, while the large majority of WT subunits exist in channels containing two chimeric subunits (Fig 3b). This indicates that the presence of a single WT subunit is sufficient for chemical activation of the channel with Yoda1. This observation is consistent with the fitting results from Fig 3a.

Next, we further performed a global fit of the dose-responses obtained for different mixtures and for WT mPiezo1. Based on equations (8) and (11), the fluorescence signal from a heterogeneous population of channel species with i number of WT subunit(s) will be a function of the saturation fraction multiplied by the total relative fraction of channel species contributing to the signal (i.e. having one or more WT subunit):

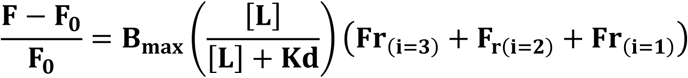

Since
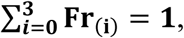
the equation can be simplified:

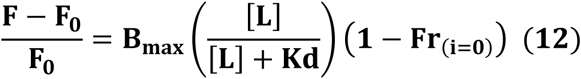

Where Fr_(i=0)_ correspond to the non-contributing fraction of channels containing three chimeric subunits. Equation (12) was used for a global fit of the data shown in Fig 3c. The fitting was done with only two shared fitted parameters, B_max_ and K_d_, while the fractions Fr_0_ were calculated for each tested plasmid mixture. The global fit produced an overall coefficient of determination R^2^ of 0.907, a B_max_ value of 4.219 ± 0.325 and a K_d_ of 9.624 ± 1.412 μM. This fitted K_d_ value is similar to the K_d_ value obtained by fitting the WT dose-response with the binding model corresponding to equation (8) (11.990 ± 1.941 μM, see Fig 3 and Table 1). Differences in the fitted Bmax can be due to fluctuations in background fluorescence q between experiments as indicated by equations (1) and (2).

Another way of confirming that each channel containing at least one WT subunit is able to open in response to Yoda1 binding is to compare the observed vs. predicted contributing fractions of Yoda1-sensitive hybrid channels. From equation (12) we can see that:

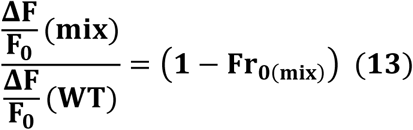

With ΔF/F_0(mix)_ the fluorescence signal observed when mixing WT and chimeric subunits, ΔF/F_0(WT)_ the fluorescence signal observed for WT channels and 1-Fr_0(mix)_ the observed contributing fraction of channels obtained when mixing subunits in a given ratio. We determined the observed contributing fraction (1-Fr_0(mix)_ by calculating the term on the left of equation (13) for each Ch:WT ratio. This number was averaged from six Yoda1 concentrations ranging between 2 μM and 75 μM. The observed contributing fractions were then plotted against the predicted contributing fractions of hybrid channels containing more than one (1 ≤ i ≤ 3), more than 2 (2 ≤ i ≤ 3) or 3 WT subunits (i = 3) using equation (11). The plots shows that the observed fractions nearly perfectly match the predicted fractions only if hybrid channels containing one or more WT subunits contribute to the fluorescence signal (Fig 3d, dotted magenta line).

## Discussion

In this study, we created chimeras between Piezo1 and Piezo2 to identify the minimal region required for the selective binding of the agonist Yoda1. In principle, the Yoda1 binding site on Piezo1 could be directly identified by solving the structure of the agonist-channel complex. However, the current resolution of the mPiezo1 structure (4.8Å) is too low to resolve most side chains in the protein. This would preclude the localization of the 21-atom Yoda1 molecule. On the other hand, the agonist binding site could be identified in functioning Piezo1 channels using a spectroscopic nanopositioning approach(29). However, such spectroscopic measurements would require fluorescent versions of Yoda1 which retain the same pharmacological properties.

Our data show that the strict selectivity of Yoda1 towards mPiezo1 originates from a minimal protein region spanning residues 1961 to 2063. This region contains 17 residues that are not conserved between the two mammalian Piezo isoforms and that are dispersed into three clusters (Supplementary Fig 1). The mPiezo1 channel becomes fully insensitive to Yoda1 only when all three clusters are exchanged with their counterpart from the mPiezo2 primary sequence. Any chimeric combination made by exchanging two out of three clusters yielded chimeras with some degree of Yoda1 sensitivity (Fig 2b). Thus, some, if not all, residues from each cluster are required to form the Yoda1 binding site. We do not know yet whether Yoda1 directly interacts with these residues or whether some of these residues are required to form a binding site in a nearby region. Hence, we envisage two hypotheses to explain the origin of Yoda1 isoform-selectivity. First, Yoda1 may directly interact with Piezo1-specific residues. In this case, Yoda1 must directly interact with some of the identified residues in each cluster in the region 1961-2063. Second, Yoda1 may directly interact with residues that are conserved in Piezo1 and Piezo2 but whose conformation depends on the presence of non-conserved residues in each cluster in the l961-2063 region. In this case the selectivity of Yoda1 must originate from a difference in the conformation of the binding site rather than from a difference in the chemical composition of its residues. Future studies will be needed to distinguish between these two possibilities.

Activation of the Yoda1-insensitive Ch1961-2063 chimera by negative pressure in a cell-attached patch requires higher pressures than WT mPiezo1 channels (Fig 2e). Incidentally, the threshold for mechanical activation of mPiezo2 channels is also higher than for mPiezo1(26). Hence, the region identified here as necessary for agonist binding appears as an important region to regulate the mechanical sensitivity of Piezo channels.

The predicted fractions of hybrid channels obtained by mixing WT and mutant subunits are true if only two assumptions are satisfied. The first is that the cellular expression of both subunits is proportional to the quantity of transfected plasmid. This seems to be the case according to the similar level of ionic current measured in cells transfected with the individual plasmids (Fig 2d). The other assumption is that both subunits randomly assemble to form trimeric channels. We do not know if this is the case. Protein-protein interaction experiments such as resonance energy transfer between fluorescent probes covalently linked to WT and chimeric subunits would confirm the existence of hybrid channels. However, the absolute quantification of the species in a heterogenous population of hybrid channels would be technically challenging to assess.

The fact that a single WT subunit suffices to confer Yoda1 sensitivity to the trimeric channel has profound mechanistic consequences regarding the gating process of the channel. Homo-multimeric ion channels with a central permeation pathway often undergo symmetrical concerted conformational rearrangements in all subunits to control the opening/closure of their pore. Hence, binding of Yoda1 to one subunit may induce a concerted transition in all subunits in the pore region that stabilizes the channel’s open state. On the other hand, Yoda1 binding in a single subunit could induce partial opening of the pore. In this case, incremental binding of the ligand in each subunit would yield distinct open states with incremental sub-conductance levels. These hypotheses could be tested by recording single channel currents from concatenated Piezo channels made by fusing a known number of WT and chimeric subunits in the same open reading frame. However, the large size of a single mammalian Piezo subunit (2400-2600 amino acids) would make those experiments challenging.

In summary, we have identified the minimal binding region of the only known isoform selective modulator of the mechanosensitive Piezo ion channel family. We further show that the binding of the agonist to only one subunit enables ligand-induced activation of the trimeric channel. This study shed lights on the gating mechanisms of Piezo proteins and will pave the way for the rationale design of small molecules with relevant pharmacological properties.

## Materials and Methods

### Molecular Cloning

A pCDNA3 plasmid containing mPiezo1 was obtained from Dr. Mikhail Shapiro (Caltech) and was originally a gift from Dr. Patapoutian (The Scripps Research Institute). A Sport6 plasmid encoding mPiezo2 was directly obtained from Dr. Pataouptian. To create the polycistronic vector pCDNA3-mPZ1-IRES-GCaMP6m, we PCR-amplified the pCDNA-mPZ1 plasmid, the internal ribosome entry site (IRES) cassette from a pIRES-eGFP plasmid and the GCaMP6m cDNA from a pGP-CMV-GCaMP6m plasmid (Addgene plasmid #40754) and assembled them using the NEBuilder HiFi DNA Assembly kit (New England Biolabs). A similar procedure was used to create pCDNA3-mPZ2-IRES-GCaMP6m, pCDNA3-GCaMP6m and the empty control vector pCDNA3. All mPZ1-mPZ2 chimeras with or withouth the IRES-GCaMP6m cassette were created using the same approach. All constructs were verified by automated Sanger sequencing (Genewiz).

### Cell culture and transfection

HEK293T cells (a gift from Dr. Mikhail Shapiro and originally purchased directly from ATCC) were cultured in standard conditions (37°C, 5% CO_2_) in a DMEM medium supplemented with Penicillin (100 U/mL), streptomycin (0.1 mg/mL), 10% sterile Fetal Bovine Serum and 1X MEM non-essential amino-acid without L-glutamine. All cell culture products were purchased from Sigma-Aldrich. Transfection was done on cells with a passage number lower than 25 using polyethylenimine (PEI, Polysciences #23966). Briefly, a sterile mixture of DNA:PEI (1:4 w/w) was prepared using sterile 100mM NaCl solution and added directly to the cell’s culture medium (220ng of total DNA was added per cm^2^ of cultured surface). Culture medium was changed 16-20 hours after transfection. For fluorescence experiments, cells seeded on clear-bottom black 96-well plates (Nunc) were transfected at ~50% confluence 2-4 days before the experiment. For electrophysiology experiments, cells seeded on uncoated coverslips were transfected at ~10% confluence 1 day prior recordings.

### Calcium imaging, osmotic shocks and chemical treatment

Fluo-8 AM was purchased from abcam (ab142773), dissolved in dimethyl sulfoxide (DMSO) and stored in 5mM aliquots at - 20°C. Yoda1 was purchased from Sigma-Aldrich (#SML1558), dissolved in DMSO and stored in 10mM aliquots at −20°C. For experiments with Fluo8-AM, cells were washed with a normal physiological solution (NPSS) containing 140mM NaCl, 5mM KCl, 2mM MgCl_2_, 1mM CaCl_2_, 10mM HEPES (pH 7.4 with HCl or NaOH) and 10mM Glucose and incubated for 1 hour in a 37°C/5% CO_2_ incubator with a NPSS solution containing 3-5 μM Fluo-8 AM. After incubation, cells were washed again with 100μL NPSS and placed on a Nikon inverted fluorescence microscope. Epifluorescence excitation was provided by a 100W mercury lamp through a 20X objective and fluorescence images were obtained using a standard GFP filter set and acquired at 1 frame/sec using a Nikon Digital Sight camera and the Nikon Digital Element D software. During recordings, 100μL of a 2X Yoda1 NPSS solution was added to the cells at t = 30 s or sometimes at t =10 s. Total DMSO concentration was kept below 1% for all tested Yoda concentrations. For osmotic shocks, 200μL of a hypotonic solution was added to the 100μL isotonic NPSS during the recording. Hypotonic solution was similar to NPSS except the NaCl concentration was reduced from 140mM to 5mM. For experiments with GCaMP6m, fluorescence imaging was done similarly except the 1 hour Fluo-8 AM treatment was replaced by a 1 hour NPSS incubation. Fluorescence images were analyzed with ImageJ.

### Cell-attached patch-clamp recordings

During recordings, the membrane potential was zeroed using a depolarizing bath solution containing 140 mM KCl, 1mM MgCl_2_, 10 mM glucose and 10 mM HEPES pH 7.3 (with KOH). Fire-polished patch pipettes with a diameter of 1-5 μm and resistance of 1-5 MΩ were filled with a recording solution containing 130 mM NaCl, 5 mM KCl, 1 mM CaCl_2_, 1 mM MgCl_2_, 10 mM TEA-Cl and 10 mM HEPES pH 7.3 (with NaOH). Stretch-activated currents were recorded in the cell-attached configuration after seal formation using a Multiclamp 700B amplifier (Axon) and a high-speed pressure clamp (HSPC-1, ALA Scientific Instruments). The membrane potential inside the patch was held at −80mV. Data were recorded at a sampling frequency of 10 kHz and filtered offline at 2 kHz using pClamp (Axon). Although the data shown are unaltered traces, peak current amplitudes were determined by adjusting the baseline at the end of the pressure pulse. No mechanosensitive ionic currents could be detected in these conditions in untransfected HEK293T cells (n = 25, data not shown).

### Statistics

For imaging, each experiment was done in duplicate and repeated n times (n values are indicated in the figures). For each experiment, the fluorescence intensity taken from 30 cells (15 cells per image stack in duplicate) was averaged. Cells were selected randomly for analysis at the beginning of the movie before Yoda1 application. Dead cells with abnormal shape or unusual high fluorescence (i.e. high cytosolic [Ca^2+^]) were excluded from analyses. The final F_(t=1min)^-^_ F_0_/F_0_ values and standard errors were calculated by averaging the mean values from n experiment. For electrophysiology, the relative peak current values and standard error (Fig 2e) are from averaging one series of recordings from n transfected cells. To test statistical difference between two mean values, we performed 2-tails unpaired Student’s t tests. The exact t-values and standard p-value ranges are indicated in each figure. The degree of freedom equals n-1. All fitting were done using OriginPro 2017.

## Acknowledgment

We thank Dr. Xiaoning Bi for lending us an inverted fluorescence microscope, Dr. Mikhail Shapiro from lending us a Multiclamp 700B amplifier and Dr. Ardem Patapoutian for the gift of the mPiezo cDNAs. We also thank David Kent, John T. Fly and Brennan Kidder for their technical help with molecular biology.

---

**Video1: Wide-field live-cell fluorescence imaging showing Yoda1-mediated Piezo1 activation**. HEK293T cells co-expressing GCaMP6m and mPiezo1 were exposed to 100 μM Yoda1 at t = 10 sec. Individual images showing GCaMP6m fluorescence were acquired at 1 frame/ sec and displayed in the movie at 5 frames/sec.

**Video2: Piezo1-dependent Yoda1-induced GCaMP6m fluorescence changes.** HEK293T cells expressing GCaMP6m only were exposed to 100 μM Yoda1 at t = 10 sec. Individual images showing GCaMP6m fluorescence were acquired at 1 frame/ sec and displayed in the movie at 5 frames/sec.

**Supplementary Figure 1: Sequence alignment (T-coffee) of the C-terminal region of mouse Piezo1 (mPZ1), Piezo2 (mPZ2) and human Piezo1 (hPZ1) and Piezo2 (hPZ2)**. The residues numbers correspond to positions used to create the chimeras. Asterisks indicate residues within the 1961-2063 region that are not conserved between mPiezo1 and mPiezo2.

